# Genomic profiling of plasma circulating tumor DNA reveals genetics and residual disease in extranodal NK/T-cell lymphoma

**DOI:** 10.1101/800409

**Authors:** Qiong Li, Wei Zhang, Jiali Li, Jingkang Xiong, Jia Liu, Ting Chen, Qin Wen, Yunjing Zeng, Li Gao, Lei Gao, Cheng Zhang, Peiyan Kong, Yao Liu, Xi Zhang, Jun Rao

**Affiliations:** Medical Center of Hematology, Xinqiao Hospital, Army Medical University, Chongqing, China; State Key Laboratory of Trauma, Burns and Combined Injury, Army Medical University, Chongqing 400037, China

**Author notes:** **Authors email** Qiong Li, Wei Zhang, Jiali Li, Jingkang Xiong, Jia Liu, Ting Chen, Qin Wen, Yunjing Zeng, Li Gao, Lei Gao, Cheng Zhang, Peiyan Kong, Yao Liu, Xi Zhang, Jun Rao. These authors contributed equally to this study (Qiong Li, Wei Zhang). Corresponding authors: Yao Liu, M. D; Xi Zhang, M. D, Ph. D; Jun Rao, M. D, Ph. D.

**Keywords:** ENKTL, circulating tumor DNA, mutation allele frequency, minimal residual disease, prognosis.

## Abstract

**Background:** Extranodal NK/T-cell lymphoma, nasal type (ENTKL), is an aggressive hematological malignancy with poor prognosis. Early detection of tumors at initial diagnosis or during routine surveillance is important for improving survival outcomes. Molecular profiling of circulating tumor DNA (ctDNA) is a promising noninvasive tool for monitoring disease status. Here, we investigated the feasible of ctDNA detection in ENTKL.

**Methods:** Plasma ctDNA was assessment were based on blood specimens that were collected from 65 patients recently diagnosed with ENKTL at the hematology medical center of Xinqiao Hospital, longitudinal samples collected under chemotherapy also included. Gene mutation spectrum of ENKTL was analyzed via cancer personalized profiling sequencing (CAPP-Seq). This study is registered with ClinicalTrials.gov (ChiCTR1800014813)

**Results:** From February 2017 to September 2019, 65 patients were enrolled, we found that the most frequently mutated genes were *KMT2D* (23.1%), *APC* (12.3%), *ATM* (10.8%), *ASXL3* (9.2%), *JAK3* (9.2%), *SETD2* (9.2%), *TP53* (9.2%), *NOTCH1* (7.7%). The mutation frequencies of *KMT2D* was significantly higher in stage III-IV, and mutations in *KMT2D, ASXL3 and JAK3* were significantly correlated with the metabolic tumor burden of the patients. Compared with tumor tissue DNA, ctDNA profiling showed good concordance. Serial ctDNA analysis showed that treatment with chemotherapy could decrease the number and mutation allele frequency of genes. Compared with PET/CT, ctDNA has more advantages for tracking residual disease in patients. In addition, we also found that mutated *KMT2D* predicted poor prognosis in patients.

**Conclusion:** Collectively, our results provide evidence that ctDNA may serve as a novel precision medicine biomarker in ENKTL.

## Introduction

Extranodal NK/T-cell lymphoma, nasal type (ENTKL), is an aggressive extranodal lymphoma of NK-cell or T-cell lineage, which is highly aggressive and has heterogenicity, with predominance in males(1–3). Current treatment strategies (such as combination chemotherapy and radiotherapy and targeted therapy) can improve the complete remission of patients, but most of them will ultimately relapse and progress(4, 5). For these patients, standard tests, including computed tomography (CT) and serum protein markers at initial diagnosis and during routine surveillance, are pivotal for therapeutic evaluation; however, these tests have limitations with low sensitivity and specificity or false positivity. Therefore, there is an urgent need for a highly sensitive, standardized and noninvasive assay that can be used for detection of early relapse and/or progression of disease.

Current treatment response criteria for non-Hodgkin lymphoma (non-HL) rely on CT scans or positron emission tomography (PET) scans. Imaging scans at initial diagnosis or recurrence can provide macro overviews of tumor volume and location, but CT scans have some limitations due to cost and radiation exposure, and PET lacks the specificity(6–8). Moreover, imaging scans cannot monitor dynamic tumor response and the clonal evolution process. Therefore, clinically validated technology is needed to overcome the current monitoring treatment response. One potential biomarker is the detection of circulating cell-free DNA (cfDNA), which comes from dying cells that release DNA fragments into the circulation, and in tumor patients, some of the cfDNA primarily originates from apoptotic and necrotic cancer cells, which carry tumor-specific alterations (named circulating tumor DNA, ctDNA)(9–12). ctDNA sequencing in plasma is a promising tool for real-time monitoring of tumor progression, and its application has been explored in multiple solid tumors. ctDNA can reflect the temporal evolution of tumors, so current studies have shown that ctDNA detection can be used to monitor minimal residual disease and track recurrence after treatment(13, 14). ctDNA sequencing is a noninvasive and tumor-specific assay, and ctDNA in circulation has a half-life of less than two hours, so ctDNA is more sensitive than protein biomarkers and more specific than routine surveillance imaging with CT scans or PET scans(15). Thus, ctDNA might become an important marker in the molecular diagnosis of tumors.

Clinical application of ctDNA has been performed in multiple types of lymphoma, such as diffuse large B-cell lymphoma (DLBCL), classical HL, follicular lymphoma and peripheral T-cell lymphoma(16–20). ctDNA assessment is a prognostic tool used prior to treatment that acts as a dynamic marker of burden during treatment by using immunoglobulin next-generation sequencing (Ig-NGS), T-cell receptor gene next-generation sequencing (TCR-NGS) and cancer personalized profiling sequencing (CAPP-Seq). Recent evidence of ctDNA assessment in peripheral T-cell lymphoma by TCR-NGS suggested that ctDNA detection in T-lineage lymphoma is feasible(21). In the present study, we thoroughly determined the molecular spectrum of ctDNA mutations in ENTKL and characterized the correlation between mutations in ctDNA and clinical factors. Paired tumor tissues and peripheral blood samples were collected to compare the concordance of tumor DNA and ctDNA. Moreover, dynamic ctDNA mutation alterations during the treatment were also observed.

## Results

### Patient characteristics

To examine the feasibility of ctDNA detection in plasma using CAPP-Seq, from February 2017 to December 2019, 65 newly diagnosed ENKTL patients were recruited, and initial and longitudinal plasma specimens were obtained from patients undergoing chemotherapy (Fig. 1). The median age at first blood draw was 45.2 years (range, 40 to 55 years), and 45 patients were men. At disease onset, 26 patients (40%) exhibited B symptoms. According to the Ann Arbor staging criteria, 14 patients were identified as stage I, 20 patients were identified as stage II, 13 patients were identified as stage III, and 18 patients were identified as stage IV (Table 1). ctDNA was successfully extracted from all patients and healthy individuals, and the average concentration of ctDNA was 3.79 hEG/ml (range, 0 to 5.39 hEG/ml).

**Figure 1.**
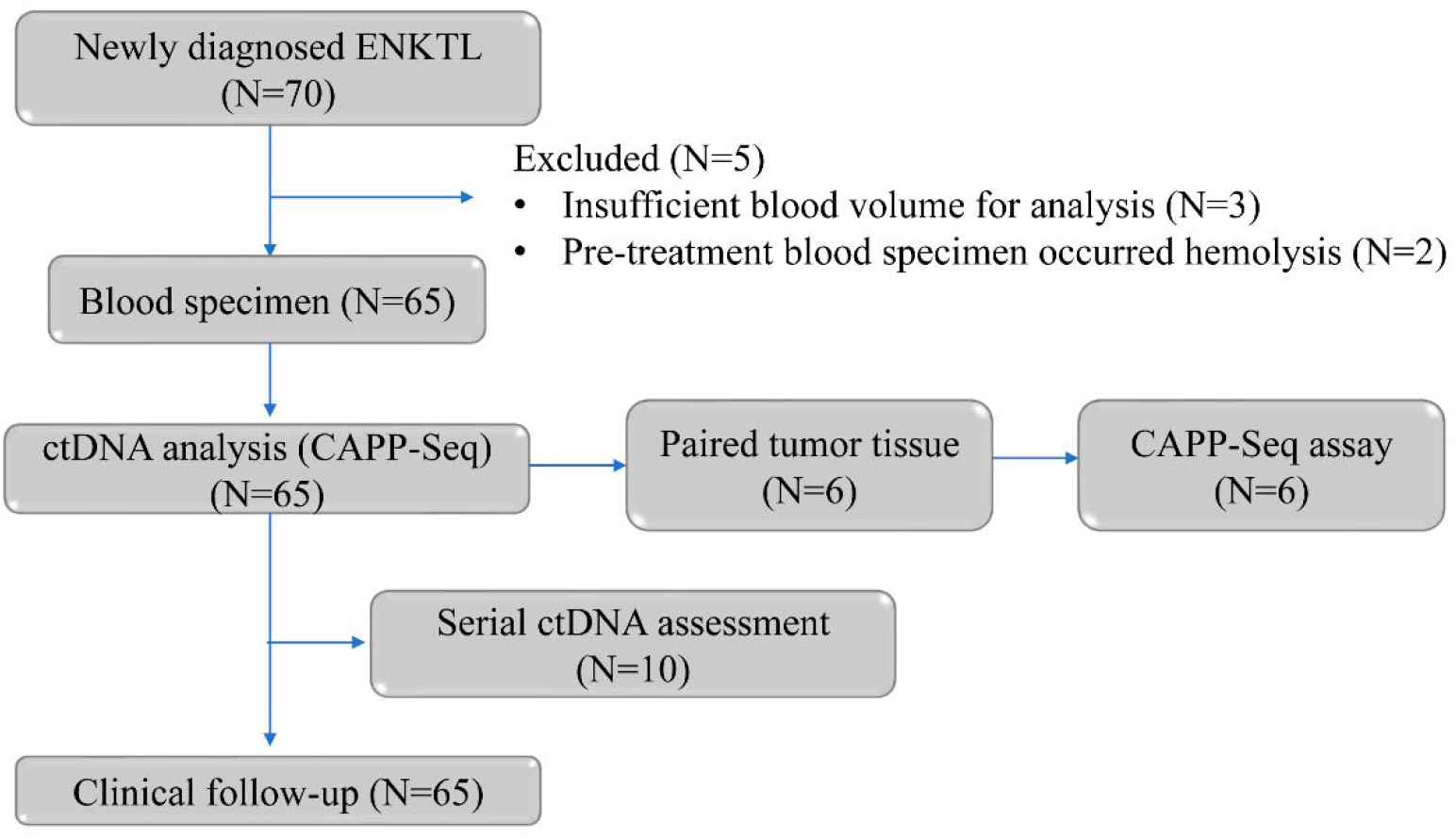
consort diagram of patient enrolment, specimen collection.

**Table 1.** Patients’ demographic and clinical characteristics

| Prognostic variables | No. | Proportion (%) |
| --- | --- | --- |
| **Gender** |  |  |
| Male | 45 | 69.23 |
| Female | 20 | 30.77 |
| **clinical stage** |  |  |
| I | 14 | 21.54 |
| II | 20 | 30.77 |
| III | 13 | 20.00 |
| IV | 18 | 27.69 |
| **B symptoms** |  |  |
| with | 26 | 40.00 |
| without | 39 | 60.00 |
| **IPI Scores** |  |  |
| 1 | 22 | 33.85 |
| 2 | 24 | 36.92 |
| 3 | 14 | 21.54 |
| 4 | 5 | 7.69 |
| **Recurrence status** |  |  |
| with | 9 | 13.85 |
| without | 56 | 86.15 |
| **Ki67 index** |  |  |
| Low | 33 | 50.77 |
| High | 32 | 49.23 |

### ctDNA mutation spectrum of newly diagnosed ENKTL

Of the 65 patients recruited in this study, in order to excluded hematopoietic clonal mutations, somatic mutations were observed in both plasma and PBMC were excluded, mutation were detected in the plasma of 49 (75.3%) patients with a median of 1.7 per sample (range, 0 to 19) (Fig. 2, upper panel). The range of mutant allele frequencies in each gene of the sample is shown in Table S1. The most frequently mutated genes were *KMT2D* (23.1%), *APC* (12.3%), *ATM* (10.8%), *ASXL3* (9.2%), *JAK3* (9.2%), *SETD2* (9.2%), *TP53* (9.2%), *NOTCH1* (7.7%) (Fig. 2, middle panel). Consistent with the somatic SNV spectrum in other tumors, we found that C>T/G>A was a preferred alteration (Fig. 2, lower panel).

**Figure 2.**
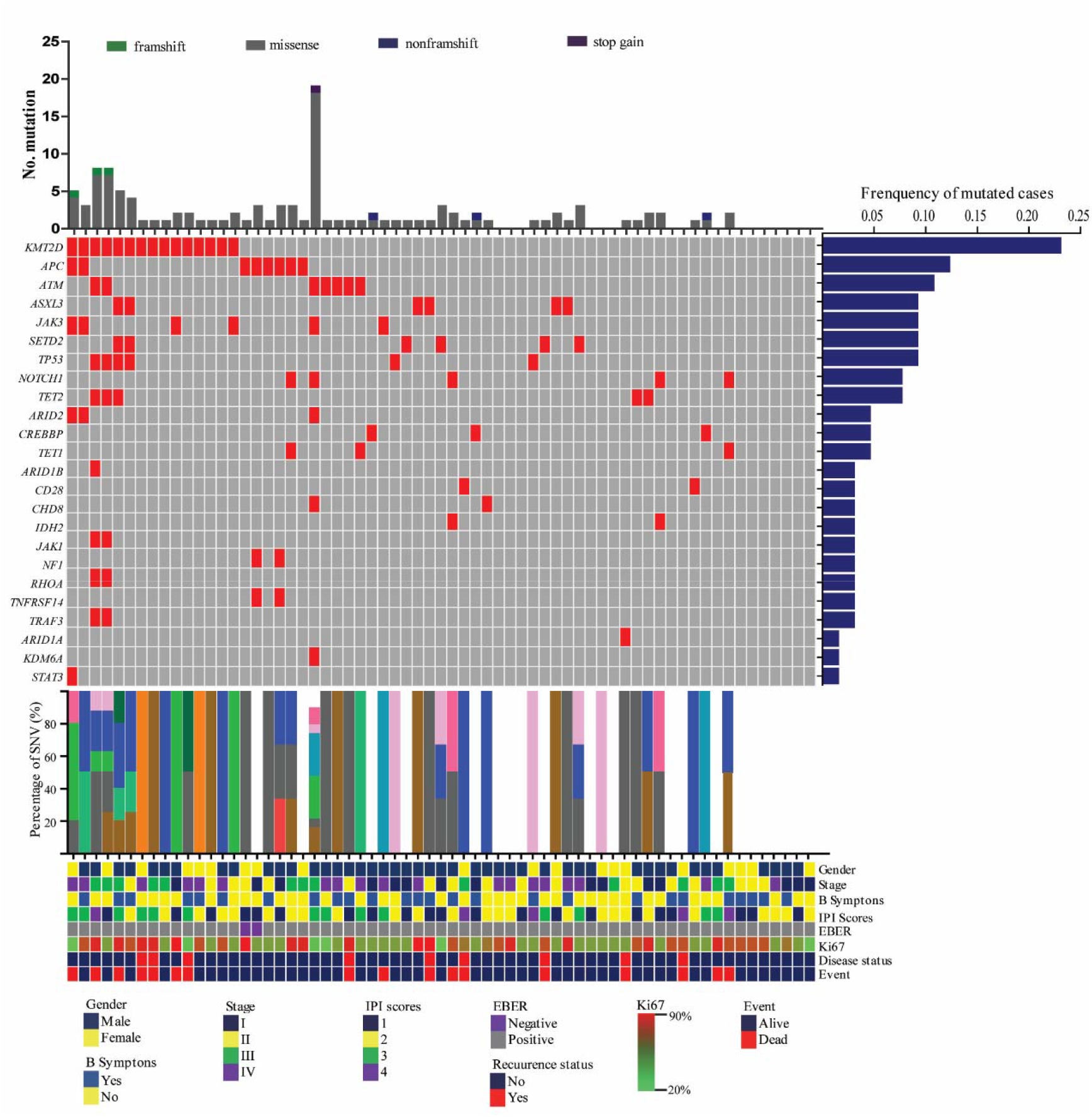
The mutational profile of newly diagnosed ENKTL. The heatmap shows individual nonsynonymous somatic mutations detected in ctDNA of newly diagnosed patients (n=65). Each row represents a gene, and each column represents a primary tumor. Mutations are color coded in red. The upper bar graph shows the number and type of mutated genes, the horizontal bar graph shows the gene mutation frequency, the middle bar graph shows the percentage of nonsynonymous somatic mutations, and the lower graph shows the clinical characteristics of each sample.

### Correlation of detectable ctDNA with clinical characteristics of ENKTL patients

To assess the concordance of the ctDNA test with the tissue NGS test, 6 paired tumor biopsies were genotyped. Compared with the results of the plasma ctDNA spectrum, most of the genes overlapped (Fig. 3A). In 4 patients, biopsy-confirmed tumor mutations were all detectable in ctDNA samples (Fig. 3B), but in patient #10, *TRAF3* variation was not detected in tumor DNA. In patient #8, *STAT3* could not be detected in tumor ctDNA (Fig. 3C). These differences might be caused by tumor anatomical heterogeneity or tumor-associated stromal tissue infiltration.

**Figure 3.**
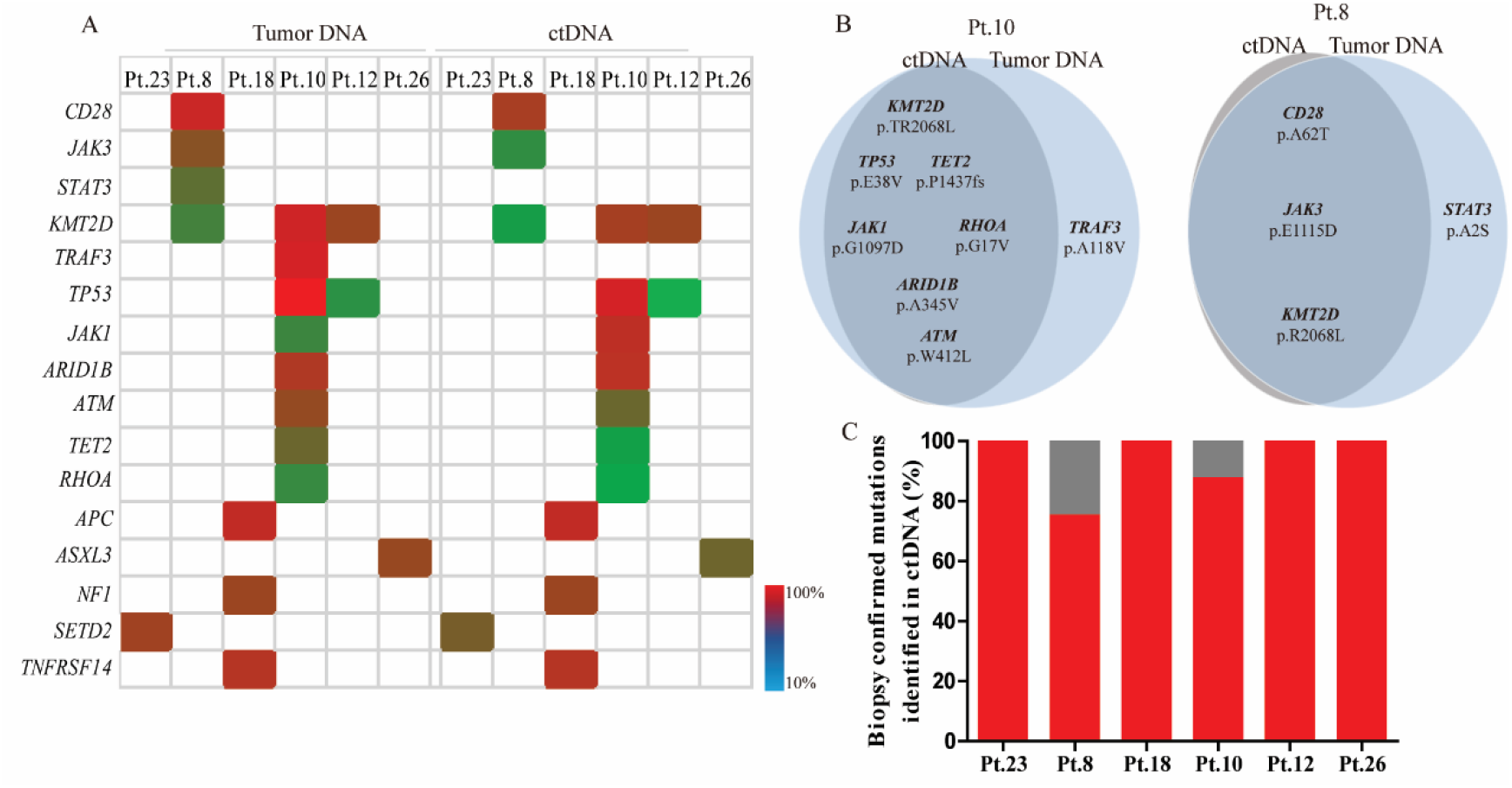
Concordance of ctDNA assessment with tissue NGS test. A. The consistent gene mutation spectrum and mutation allele frequency detected in plasma ctDNA and tumor tissue of the patients. B. Venn diagram summarizing the detailed mutations discovered in both plasma ctDNA (gray) and tumor DNA (blue). C. For each patient, the fraction of tumor biopsy-confirmed mutations that were detected in plasma ctDNA is shown. The gray portion of the bar marks the part of tumor biopsy-confirmed mutation that was not found in plasma ctDNA.

To validate the potential clinical utility of mutations detected in plasma ctDNA, we investigated the association between clinical factors and ctDNA levels. As MTV measured from PET/CT could quantitatively reflect tumor burden, we found that the plasma cfDNA concentration was significantly correlated with MTV (P=0.04) (Fig. 4A), suggesting that ctDNA could be identified as a promising biomarker for tumor stage and burden. In patients with stage III-IV disease, the mutation frequencies of *KMT2D* (11/33), *ATM* (5/33), *JAK3* (3/33) were higher than that of patients with stage I-II disease (Fig. 4B). Additionally, the mutation allele frequencies of *KMT2D, APC* and *ASXL3* were not correlated with patient stage, but the mutation frequency of *ATM, JAK3* showed significant differences (Fig. 4C). Furthermore, the correlation of the mutation status of each gene and metabolic tumor burden was investigated. We found that patients with mutations in *KMT2D, ASXL3,* and *JAK3* showed significantly higher metabolic tumor burden (Fig. 4D), suggesting that these genes might be positively correlated with disease malignancies. Recently, plasma EBV-DNA can serve as a valuable biomarker of tumor load and prognostic factor, we also evaluated the correlation between plasma EBV-DNA and clinical characteristics, we found that pre-treatment plasma EBV-DNA was positively correlated with MTV (Fig. 4E), but the correlation of EBV-DNA levels and ctDNA concentration was not significant (Fig. 4F).

**Figure 4.**
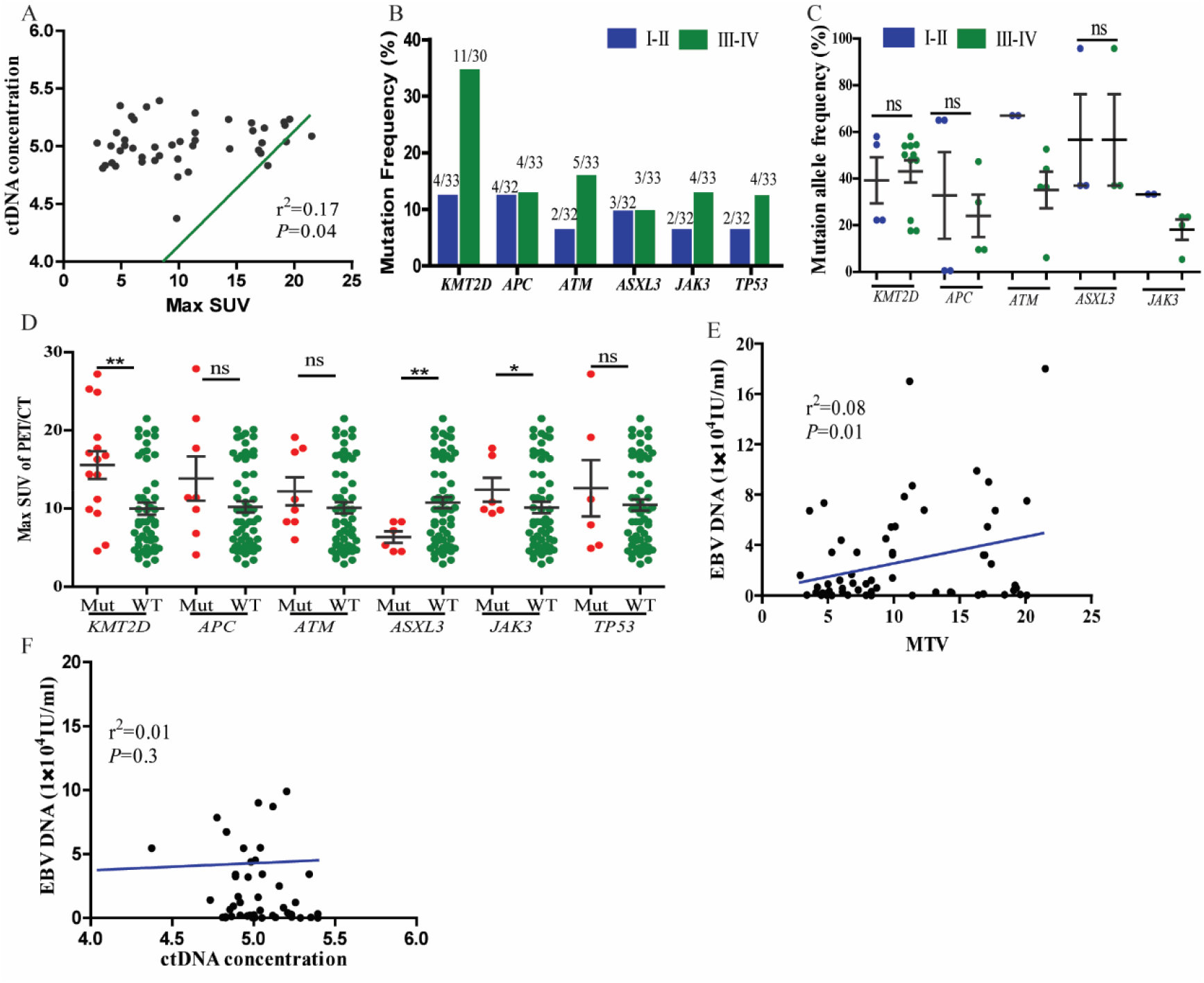
Correlation of ctDNA assessment with patient clinical characteristics. A. Linear regression of plasma ctDNA concentration with metabolic tumor volume. B. The mutation frequency of *KMT2D, APC, ATM, ASXL3, JAK3* and *TP53* in patients with different tumor stages. C. The mutation allele frequency of *KMT2D, APC, ATM, ASXL3 and JAK3* in patients with different tumor stages. D. The general metabolic tumor volume in patients with different mutation statuses of *KMT2D, APC, ATM, ASXL3, JAK3* and *TP53*. E. The correlation of plasma EBV-DNA levels and metabolic tumor volume. F. The correlation of EBV-DNA levels and ctDNA concentration.

### serial ctDNA detection during therapy could complement the response assessment of the patients, and patients with mutated KMT2D and ATM predicted poor prognosis

We further investigated the potential role of plasma ctDNA in the therapeutic monitoring of patients with chemotherapy. Ten patients were recruited, sequential plasma samples at disease diagnosed, before treatment cycle 4 (C4), and before cycle 8 (C8) were collected. 7 patients could achieve CR at C4, but in these patients, only 2 couldn’t detect mutation in plasma, suggesting that ctDNA assessment was more sensitivity than PET/CT scan, all patients couldn’t detect plasma EBV-DNA. In addition, with more cycles of chemotherapy, all patients could achieve complete molecular remission (Fig. 5). In these 10 patients, we also observed one relapsed, and the relapsed spectrum was similar with the original spectrum.

**Figure 5.**
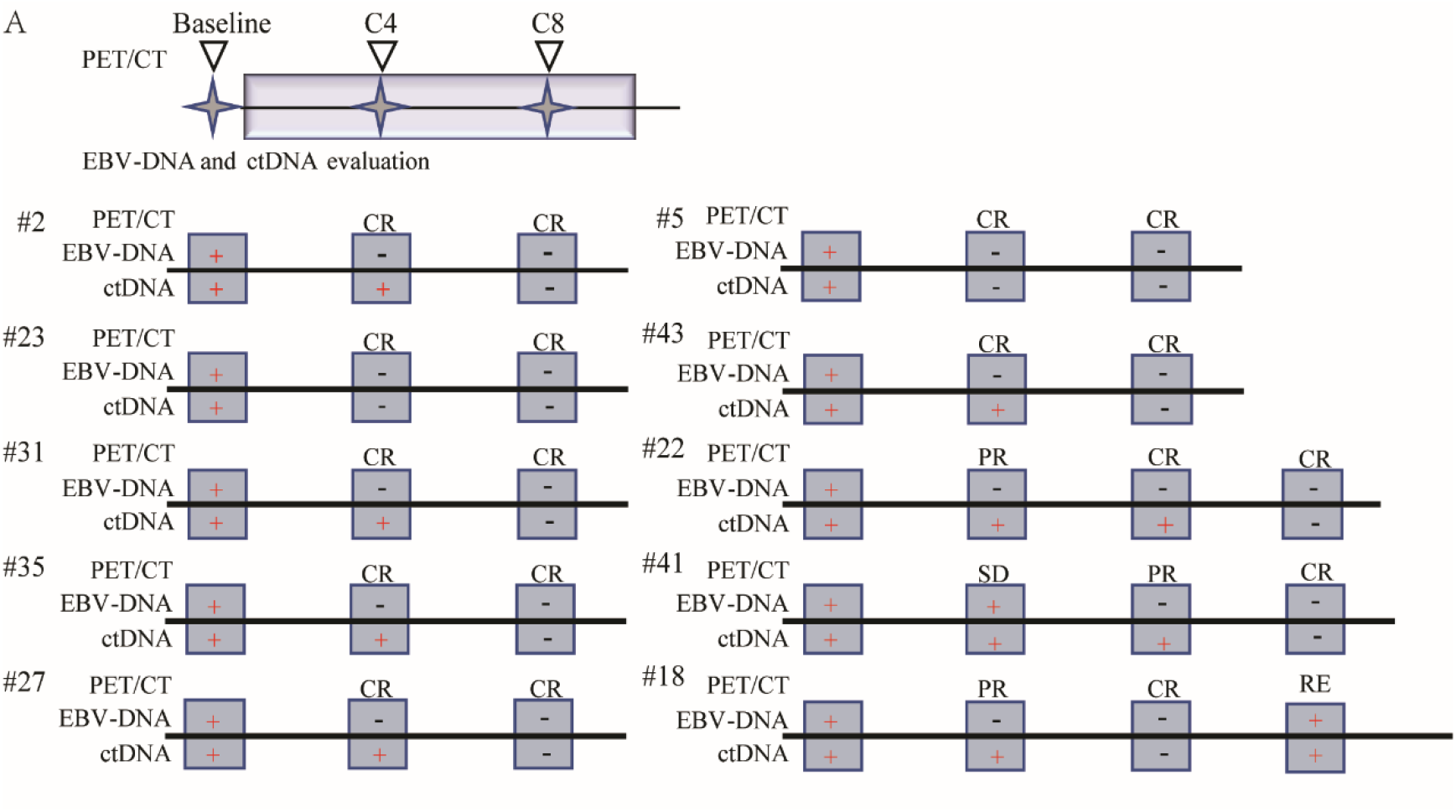
Serial ctDNA detection during therapy could complement the response assessment of the patients. Treatment response assessment using PET/CT is combined with genomic testing of ctDNA and plasma EBV-DNA levels. Plasma and imaging were performed at baseline prior to treatment initiation, before start of cycle 4 (C4), and before cycle 8 (C8). Displayed are the patients recruited with serial plasma collection and their response to study treatment using plasma EBV-DNA, ctDNA and PET/CT.

Because KMT2D was the most frequent somatic mutations, the expression of KMT2D was investigated, we found that the KMT2D expression of patients with mutated was higher than that of patients with wild type (Fig. 6A). Next, the prognosis value of plasma ctDNA mutation status and EBV-DNA level was also investigated. Kaplan-Meier analysis estimated that patients with mutated *KMT2D* and *ATM* had a shorter OS (Fig. 6B, 6C), No significant difference was observed in patients with other mutated genes (Fig. 6D-G), which might be due to the small sample recruited in this clinical trial. In addition, patients with high EBV-DNA level showed significantly poor prognosis (Fig. 6H). The results of univariate and multivariate analysis for risk factor of OS in patients were summarized in Tables 2. Univariate analysis of factors revealed that stage, IPI scores, recurrence status, EBV-DNA levels, *KMT2D* mutation and *ATM* mutation were independent prognostic indicators of the overall survival of patients, and multivariate analysis showed that EBV-DNA levels, *KMT2D* mutation and *ATM* mutation was solely prognostic factor. All these results suggested that dynamic detection of ctDNA during therapy could better reflect disease status compared with PET/CT scan, and patients with mutated *KMT2D,* and *ATM* predicted poor prognosis.

**Table 2.** Univariate and multivariate analyses of the overall survival of patients

| Prognostic variables | Univariate analysis |  | Multivariate analysis |  |
| --- | --- | --- | --- | --- |
|  | HR (95% CI) | *p value* | HR (95% CI) | *p value* |
| Gender (female vs. male) | 0.712(0.279-1.812) | 0.476 |  |  |
| Stage (I-II vs. III-IV) | 2.625(1.043-6.604) | 0.04 | 1.159(0.253-5.311) | 0.849 |
| B symptoms (with vs. without) | 1.234(0.512-2.973) | 0.64 |  |  |
| IPI | 2.595(1.073-6.277) | 0.034 | 0.565(0.118-2.712) | 0.476 |
| Recurrence status | 3.346(1.364-8.204) | 0.008 | 2.705(0.941-7.776) | 0.065 |
| MTV | 2.069(0.843-5.078) | 0.112 |  |  |
| ctDNA concentraion | 2.975(0.984-8.997) | 0.053 |  |  |
| EBV-DNA level | 3.672(1.396-9.664) | 0.008 | 3.174(1.091-9.231) | 0.034 |
| KMT2D mutation (without vs. with) | 0.289(0.119-0.702) | 0.006 | 0.266(0.086-0.828) | 0.022 |
| APC mutation (without vs. with) | 0.550(0.16-1.895) | 0.344 |  |  |
| ATM mutation (without vs. with) | 0.271(0.098-0.752) | 0.012 | 0.212(0.064-0.700) | 0.011 |
| ASXL3 mutation (without vs. with) | 2,613(0.347-19.68) | 0.351 |  |  |
| JAK3 mutation (without vs. with) | 0.569(0.166-1.949) | 0.369 |  |  |
| SETD2 mutation (without vs. with) | 0.804(0.184-3.507) | 0.771 |  |  |
| TP53 mutation (without vs. with) | 0.615(0.179-2.115) | 0.441 |  |  |
| NOTCH1 mutation (without vs. with) | 0.410(0.120-1.406) | 0.156 |  |  |

**Figure 6.**
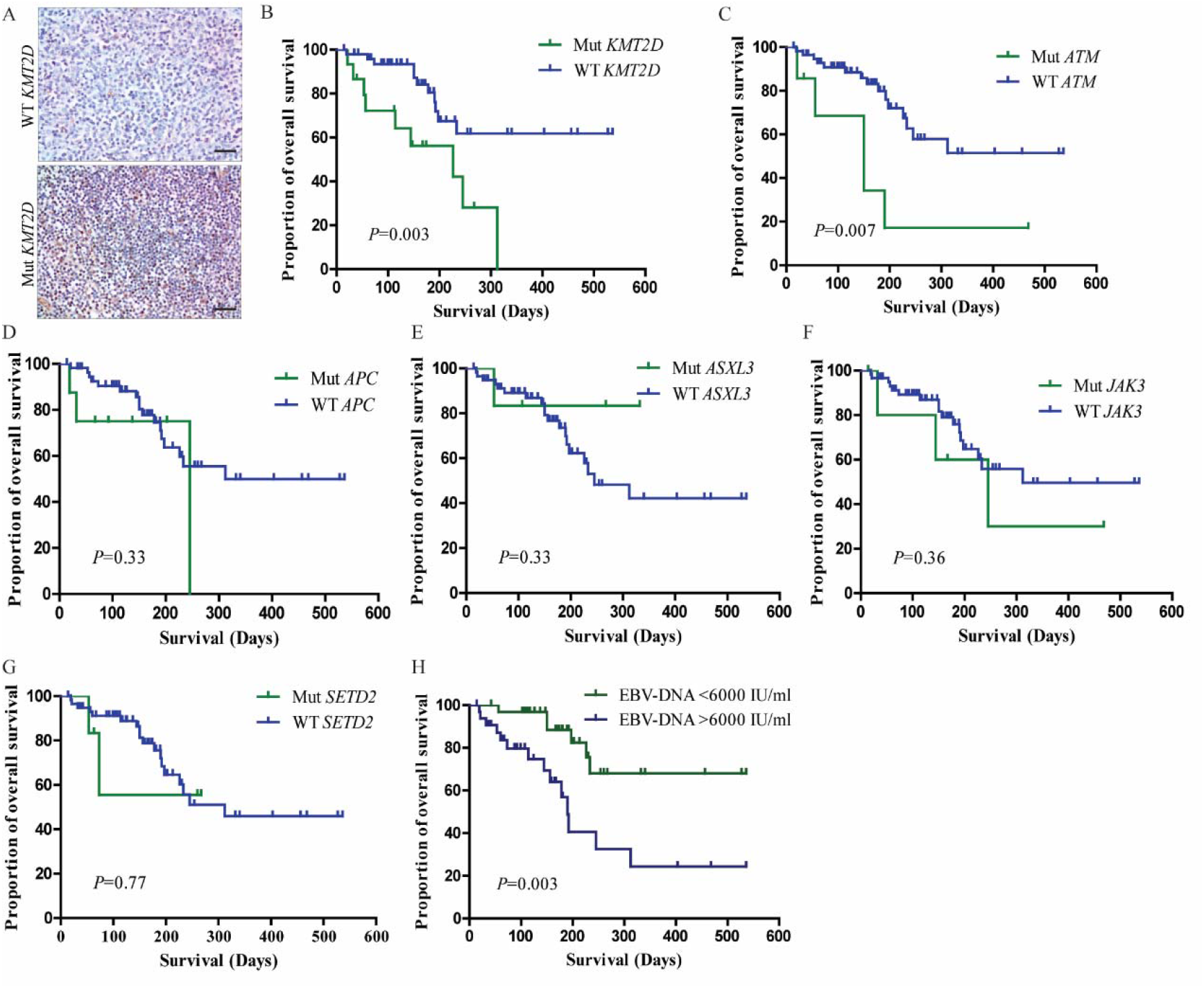
Prognostic value of frequently mutated genes on patients with ENKTCL. A. Immunohistopathology staining of KMT2D in patients with mutated KMT2D and wild type, scale bar, 25um. Kaplan-Meier estimated the OS of plasma *KMT2D* mutation status in patients with ENKTL (B), *ATM* (C), *APC* (D), *ASXL3* (E), *JAK3* (F), *SETD2* (G), plasma EBV-DNA levels (H)

## Discussion

This study was the first to determine the plasma ctDNA mutation gene spectrum of ENKTL using CAPP-Seq and demonstrated that ctDNA sequencing could be considered a promising biomarker for monitoring the minimal residual disease (MRD) of patients. The most frequently mutated genes in plasma ctDNA were *KMT2D* (23.1%), *APC* (12.3%), *ATM* (10.8%), *ASXL3* (9.2%), *JAK3* (9.2%), *SETD2* (9.2%), *TP53* (9.2%), *NOTCH1* (7.7%). ctDNA concentration was positively correlated with tumor stage and MTV. Consistent with tumor tissue, almost all of the gene mutations detected in plasma ctDNA could be found in tumor DNA, and further analysis demonstrated that a decreased number and MAF of certain genes in plasma ctDNA could complement the therapeutic response of the patients, and patients with mutated *KMT2D* and *ATM* had poor prognosis, suggesting that the gene mutations in plasma could serve as a promising biomarker for the diagnosis or monitoring of ENKTL disease courses.

Tumor biopsies can only offer information about lymphoma at one specific time in a specific area and can be accompanied with complications, and PET/CT may lead to false negatives due to technology limitations or false positive caused by tumor flares or pseudoprogression(22). ctDNA detection is an optimal noninvasive and specific technique to overcome these traditional monitoring methods. A rapidly growing body of evidence has established that ctDNA detection during curative therapies can identify patients for whom there remains evidence of residual, radiographically occult cancer. In DLBCL, ctDNA can be used to predict tumor load and treatment outcome of newly diagnosed patients, and patients with a pretreatment ctDNA level above the median had a significantly poorer prognosis than those below the median(23). Later, Kurtz et al. reported that serial ctDNA measurements of 125 patients were an effective tool for prognostication, patients with 2-log or greater decreases in ctDNA had a better event-free survival (EFS) after one cycle of therapy, and patients with 2.5-log or greater reduction in ctDNA had a significantly better EFS after two cycle treatments. These studies suggested that serial ctDNA detection could be used to assess the therapy response dynamically and guide personalized target therapy. In another study, Daigle et al. found that mutation of *CREBBP* in R/R DLBCL had the greatest association with tazemetostat treatment (an EZH2 inhibitor), and patients with mutations in *PIM1*, *BCL6*, *HIST1H1E* and *TP53* lacked response to treatment(24). In HL, Valeria et al. determined the molecular mutation spectrum using ctDNA measurement. The most mutated genes were *STAT*, *TNFAIP3* and *ITPKB*, and detection sensitivity was 87.5%, suggesting that ctDNA could reflect the spectrum of tumor tissue mutation well. Compared with newly diagnosed and refractory HL mutation profiles, most mutations overlapped, and mutations of *STAT*, *TNFAIP3, GNA13* and *ITPKB* might be an ancestral clone persisted throughout disease course(20). Although only a few studies have focused on assessing ctDNA in T-cell lymphoma, preliminary evidence has also demonstrated that ctDNA measurement in TCL is feasible(21, 25). Our study is the first to determine the mutation spectrum of ctDNA in ENKTL. The most common mutation genes are *KMT2D, APC, ATM, ASXL3, JAK3, SETD2, TP53* and *NOTCH1*. Our results revealed that ctDNA could reflect MTV, and by longitudinally profiling patients treated with chemotherapy, we provide evidence that serial ctDNA assessment could monitor the disease status, and the results could partially reflect which mutated gene was sensitive to chemotherapy drugs. However, only one of our 10 patients recruited for serial monitoring relapsed; for relapsed patients, whether primary mutation clone recurrence or new mutation occupies the main role needs to be further investigated. In addition, future trials should include more patients and long-term follow-up to explore the plasma gene clone evolution process.

Plasma cfDNA includes tumor cell-derived plasma ctDNA and normal cell apoptosis-derived cfDNA, which are DNA fragments. ctDNA could be a more comprehensive reflection of gene mutations in all tumors than conventional biopsy(26, 27). Therefore, ctDNA-based detection methods are critical for identifying which mutation is tumor-specific. As sensitivity increases with the development of new methods and because age-related somatic mutations could also lead to hematopoietic clonal expansions, the possibility of false-positive results might occur(28, 29). Recently, Razavi et al showed that clonal hematopoiesis was major contributor to plasma variants using a high-intensity sequencing assay of matched plasma DNA, tumor tissue and white blood cells, emphasizing that matched cfDNA-white blood cells was critical for screening accurate somatic mutations(30). In the present study, to decrease the bias caused by clonal hematopoiesis associated mutations, the age range of the included patients was narrow, matched ctDNA-PBMC sample was applied, and the sequencing panel was conducted according to a previous gene-expression profiling study on ENKTL. Zhao et al. confirmed that the most frequently mutated genes in NKTCL were the RNA helicase gene *DDX3X*, tumor suppressors (*TP53* and *MGA*), JAK-STAT pathway molecules (*STAT3* and *STAT5b*) and epigenetic modifiers (*MLL2, ARID1A, EP300* and *ASXL3*)(31). In addition to those recurrent mutations, *JAK3, KMT2D, EZH2, NOTCH1* and *TET2* were also identified by different groups. Combined with the most frequent gene mutation spectrum in Chinese individuals, we selected 41 mutated genes for our study. Interestingly, we found that *KMT2D* alterations possessed a higher frequency (23.1%) and which were all non-synonymous mutations (p.R2068L, p.D4378E, p.R755W, p.P2382S, p.V160L, p.L2316V). KMT2D belongs to a family of mammalian histone H3 lysine 4 (H3K4) methyltransferase, it is frequently mutated in Kabuki syndrome and various cancers, which acts as a scaffold protein within the complex depending on cell type and stage of differentiation(32). In acute myeloid leukemia, follicular lymphoma and DLBCL, KMT2D could inhibit tumorigenesis and metastasis(33, 34), but in solid cancer, such as prostate cancer, KMT2D was critical factor for promoting tumor cell proliferation(35). In our study, we found that the KMT2D expression of patients with mutated was higher than that of patients with wild type, suggesting that KMT2D might be oncogene in NKTCL lymphoma. However, the underlying significance in NKTCL has not been explored to date, and further studies regarding function of KMT2D mutation in NKTCL lymphomagenesis are urgently recommended. To our knowledge, this is the first study to prospectively evaluate the potential utility of ctDNA analysis in NKTCL patients. We have demonstrated that ctDNA assessment could predict the therapy response in NKTCL, and the highest mutation frequencies were in *KMT2D* and *APC*. Our results could define the optimal strategy for patient follow-up.

## Materials and methods

### 1. Study design and patient selection

In this prospective cohort study, 65 patients recently diagnosed with ENKTL at the hematology medical center of Xinqiao Hospital from February 2017 to December 2019 were enrolled (ClinicalTrials identifier: ChiCTR1800014813). All consecutive patients who were deemed appropriate for this study during the study period were included without selection. Histological diagnoses were established independently by at least two experienced senior pathologists according to the WHO classification of Tumors of Hematopoietic and Lymphoid tissue criteria. All patients underwent baseline staging using laboratory, radiographic, and bone marrow examinations.

Eastern Cooperative Oncology Group (ECOG) performance status was assessed at diagnosis. Stage was evaluated in accordance with the Ann Arbor staging system. The International Prognostic Index (IPI) was calculated based on serum lactate dehydrogenase, stage, extranodal status and performance status. Patient characteristics and treatment regimens of each therapy cycle were collected from each patient. Three healthy individuals were also recruited as control participants to test the accuracy of our sequencing platform for ctDNA profiling. All participants provided informed written consent before undergoing any study-related procedures in accordance with the Declaration of Helsinki. This study was approved by the China Ethics Committee of Registering Clinical Trials (ChiECRCT-20180005).

After the initial stage assessment, all patients were given front-line therapy according to National Comprehensive Cancer Network (NCCN) guidelines. Patients were reviewed routinely with a combination of clinical assessment and CT or fluorodeoxyglucose-PET (FDG-PET). FDG-PET was often used as an interim scan, and metabolic tumor volume (MTV) was determined from the initial and interim PET images using PET Edge software (MIMSoftware Inc., Cleveland, OH, USA). ctDNA profiling was conducted according to the therapy response during the patient’s therapy course.

### 2. Sample collection and DNA extraction

Before treatment, peripheral blood samples were collected using 10 ml EDTA vacutainer tubes and processed within 4 hours. cfDNA was extracted from plasma using the QIAamp Circulating Nucleic Acid Kit (Qiagen, Valencia, California) following the manufacturer’s instructions. Genomic DNA was extracted from peripheral mononuclear cells and formalin fixed paraffin-embedded (FFPE) tissue. DNA concentration and quality were estimated using a Qubit fluorometer (Invitrogen). cfDNA quality was assessed using an Agilent 2100 Bioanalyzer and DNA HS Kit (Agilent Technologies, Palo Alto, CA, USA).

### 3. Multiregional targeted NGS of patients’ plasma and tumor sample

Sequencing was performed on FFPE tumor DNA, plasma cfDNA and PBMC, and targeted sequencing gene panels including coding exons and splice sites of 41 genes (Yuanqi Biopharmaceutical Co., Shanghai, China) that are recurrently mutated in NK/T-cell lymphoma were specifically designed for this study, including ADAM3A, *APC, ARID1A, ARID1B, ARID2, ASXL3, ATM, BCOR, BCORL1, CD28, CHD8, CREBBP, DDX3X, DNMT3A, EP300, EZH2, FYN, IDH2, IL2RG, JAK1, JAK3, KDM6A, KMT2A, KMT2D, MGA, NF1, NOTCH1, PRDM1, PTPN1, RHOA, SETD2, SOCS1, STAT3, STAT5B, STAT6, TET1, TET2, TNFRSF14, TP53, TRAF3* and *ZAP608*. Tumor DNA was sheared through sonication before library construction to obtain an almost 200-bp fragment for cfDNA, which possesses a fragmented DNA nature. No additional fragmentation was performed before library construction. The NGS libraries were constructed using the SureSelect Library Prep Kit (Agilent Technologies, Palo Alto, CA, USA). Quantification of the library was performed using the Agilent DNA 1000 Kit (Agilent Technologies). Sequencing was performed following the manufacturer’s protocol on the Illumina MiSeq system (Illumina, San Diego, CA). For tumor DNA, the mean depth of each sample was 2500×. The length of cfDNA fragments primarily ranged from 100 to 200 bp, and the average coverage depths for sequencing were 813.42× (range, 462×-6513×), with an average of 5% of the target sequence being covered with sufficient depth for variant calling. Bioinformatics analysis was performed to verify the sample sequence and mutation site and calculate the mutated allele frequency (MAF) compared with the human genome sequence (hg19) using Burrows-Wheel Aligner (BWA) sequence alignment software. Samtools version 1.3 was used for single nucleotide variant (SNV)/indel calling and filter workflow.

### 4. Statistical analysis

Statistical analysis was conducted using SPSS software (Version 18.0, LEAD Corp). Descriptive statistics were used to analyze clinical, demographic and genetic test result characteristics. Correlation of gene mutation and clinicopathologic features of patients were conducted using the Pearson χ^2^ test. Overall survival (OS) measured the proportion of patients who were alive at a specific time after diagnosis. Survival estimates were obtained using the Kaplan–Meier method, and comparisons were made using a log-rank test. Unpaired Student’s t-test for two groups were used in this study. A COX proportional regression model was used to calculate the survival hazard ratio (HR). Statistical differences were considered significant if the *P* value was less than 0.05.

## List of abbreviations

ENTKL: Extranodal NK/T-cell lymphoma, nasal type
CT: Computed tomography
Non-NHL: non-Hodgkin lymphoma
cfDNA: circulating cell-free DNA
ctDNA: circulating tumor DNA
DLBCL: diffuse large B-cell lymphoma
CAPP-Seq: cancer personalized profiling sequencing
MTV: metabolic tumor volume
ADAM3A: ADAM Metallopeptidase Domain 3A
*APC*: Adenomatous Polyposis Coli Protein
*ARID1A*: AT-Rich Interaction Domain 1A
*ARID1B*: AT-Rich Interaction Domain 1B
*ARID2*: AT-Rich Interaction Domain 2
*ASXL3*: ASXL Transcriptional Regulator 3
*ATM*: Ataxia Telangiectasia Mutated
*BCOR*: BCL6 Corepressor
*BCORL1*: BCL6 Corepressor Like 1
*CHD8*: Chromodomain Helicase DNA Binding Protein 8
*CREBBP*: CREB Binding Protein
*DDX3X*: DEAD-Box Helicase 3 X-Linked
*DNMT3A*: DNA Methyltransferase 3 Alpha
*EP300*: E1A Binding Protein P300
*EZH2*: Enhancer Of Zeste 2 Polycomb Repressive Complex 2 Subunit
*FYN*: Src Family Tyrosine Kinase
*IDH2*: Isocitrate Dehydrogenase (NADP(+)) 2, Mitochondrial
*IL2RG*: Interleukin 2 Receptor Subunit Gamma
*JAK1*: Janus Kinase 1
*JAK3*: Janus Kinase3
*KDM6A*: Lysine Demethylase 6A
*KMT2A*: Lysine Methyltransferase 2A
*KMT2D*: Lysine Methyltransferase 2D
*MGA*: MAX Dimerization Protein
*NF1*: Neurofibromin 1
*NOTCH1*: Notch Receptor 1
*PRDM1*: Positive Regulatory Domain I-Binding Factor 1
*PTPN1*: Protein Tyrosine Phosphatase Non-Receptor Type 1
*RHOA*: Ras Homolog Family Member A
*SETD2*: SET Domain Containing 2
*SOCS1*: Suppressor of Cytokine Signaling 1
*STAT3*: Signal Transducer and Activator of Transcription 3
*STAT5B*: Signal Transducer and Activator of Transcription 5B
*STAT6*: Signal Transducer and Activator of Transcription 6
*TET1*: Tet Methylcytosine Dioxygenase 1
*TNFRSF14*: TNF Receptor Superfamily Member 14
*TP53*: Tumor Protein P53
*TRAF3*: TNF Receptor Associated Factor 3
*ZAP608*: Zeta Chain Of T Cell Receptor Associated Protein Kinase 608
MAF: mutated allele frequency
SNV: single nucleotide variant

## Declaration

### Ethics approval and consent to participate

The study was approved by china ethics committees of registering clinical trials (ChiECRCT-2018005)

## Consent for publication

All authors reviewed and approved the manuscript.

## Availability of data and materials

All data generated and analyzed during this study are included in this article and its supplementary information files.

## Author contributions

JR, YL, and XZ conceived and designed the study. JR, WZ and QL developed the methodology and analyzed and interpreted the data. JR and WZ wrote the manuscript. All authors reviewed and revised the manuscript.

## Acknowledgments

We thank Dr. Yimei Feng for their assistance with the statistical analysis and Qing Xu for her encouragement. The authors also thank their colleagues for helpful comments.

## Supplementary Materials

Table S1: The range of mutant allele frequencies in each gene of each sample.

## References

1. Tse E, and Kwong YL. How I treat NK/T-cell lymphomas. Blood. 2013;121(25):4997–5005.

2. Suzuki R. Pathogenesis and treatment of extranodal natural killer/T-cell lymphoma. Semin Hematol. 2014;51(1):42–51.

3. Xiong J, and Zhao W. What we should know about natural killer/T-cell lymphomas. Hematol Oncol. 2019;37 Suppl 1:75–81.

4. Kwong YL, Kim WS, Lim ST, Kim SJ, Tang T, Tse E, et al. SMILE for natural killer/T-cell lymphoma: analysis of safety and efficacy from the Asia Lymphoma Study Group. Blood. 2012;120(15):2973–80.

5. Au WY, Weisenburger DD, Intragumtornchai T, Nakamura S, Kim WS, Sng I, et al. Clinical differences between nasal and extranasal natural killer/T-cell lymphoma: a study of 136 cases from the International Peripheral T-Cell Lymphoma Project. Blood. 2009;113(17):3931–7.

6. Adams HJ, Nievelstein RA, and Kwee TC. Prognostic value of complete remission status at end-of-treatment FDG-PET in R-CHOP-treated diffuse large B-cell lymphoma: systematic review and meta-analysis. Br J Haematol. 2015;170(2):185–91.

7. Trotman J, Luminari S, Boussetta S, Versari A, Dupuis J, Tychyj C, et al. Prognostic value of PET-CT after first-line therapy in patients with follicular lymphoma: a pooled analysis of central scan review in three multicentre studies. Lancet Haematol. 2014;1(1):e17–27.

8. Kim SJ, Choi JY, Hyun SH, Ki CS, Oh D, Ahn YC, et al. Risk stratification on the basis of Deauville score on PET-CT and the presence of Epstein-Barr virus DNA after completion of primary treatment for extranodal natural killer/T-cell lymphoma, nasal type: a multicentre, retrospective analysis. Lancet Haematol. 2015;2(2):e66–74.

9. Forshew T, Murtaza M, Parkinson C, Gale D, Tsui DW, Kaper F, et al. Noninvasive identification and monitoring of cancer mutations by targeted deep sequencing of plasma DNA. Sci Transl Med. 2012;4(136):136ra68.

10. Bettegowda C, Sausen M, Leary RJ, Kinde I, Wang Y, Agrawal N, et al. Detection of circulating tumor DNA in early- and late-stage human malignancies. Sci Transl Med. 2014;6(224):224ra24.

11. Siravegna G, Marsoni S, Siena S, and Bardelli A. Integrating liquid biopsies into the management of cancer. Nat Rev Clin Oncol. 2017;14(9):531–48.

12. Crowley E, Di Nicolantonio F, Loupakis F, and Bardelli A. Liquid biopsy: monitoring cancer-genetics in the blood. Nat Rev Clin Oncol. 2013;10(8):472–84.

13. Zhou J, Huang A, and Yang XR. Liquid Biopsy and its Potential for Management of Hepatocellular Carcinoma. J Gastrointest Cancer. 2016;47(2):157–67.

14. Esposito A, Criscitiello C, Locatelli M, Milano M, and Curigliano G. Liquid biopsies for solid tumors: Understanding tumor heterogeneity and real time monitoring of early resistance to targeted therapies. Pharmacol Ther. 2016;157:120–4.

15. Chen Y, George AM, Olsson E, and Saal LH. Identification and Use of Personalized Genomic Markers for Monitoring Circulating Tumor DNA. Methods Mol Biol. 2018;1768:303–22.

16. Roschewski M, Dunleavy K, Pittaluga S, Moorhead M, Pepin F, Kong K, et al. Circulating tumour DNA and CT monitoring in patients with untreated diffuse large B-cell lymphoma: a correlative biomarker study. Lancet Oncol. 2015;16(5):541–9.

17. Scherer F, Kurtz DM, Newman AM, Stehr H, Craig AF, Esfahani MS, et al. Distinct biological subtypes and patterns of genome evolution in lymphoma revealed by circulating tumor DNA. Sci Transl Med. 2016;8(364):364ra155.

18. Suehara Y, Sakata-Yanagimoto M, Hattori K, Nanmoku T, Itoh T, Kaji D, et al. Liquid biopsy for the identification of intravascular large B-cell lymphoma. Haematologica. 2018;103(6):e241–e4.

19. Delfau-Larue MH, van der Gucht A, Dupuis J, Jais JP, Nel I, Beldi-Ferchiou A, et al. Total metabolic tumor volume, circulating tumor cells, cell-free DNA: distinct prognostic value in follicular lymphoma. Blood Adv. 2018;2(7):807–16.

20. Spina V, Bruscaggin A, Cuccaro A, Martini M, Di Trani M, Forestieri G, et al. Circulating tumor DNA reveals genetics, clonal evolution, and residual disease in classical Hodgkin lymphoma. Blood. 2018;131(22):2413–25.

21. Melani C, Pittaluga S, Yee L, Lucas A, Shovlin M, Jacob A, et al. Next-Generation Sequencing Based Monitoring of Circulating-Tumor DNA in Untreated Peripheral T-Cell Lymphoma. Blood. 2017;130(Suppl 1):2728-.

22. Cheson BD, Ansell S, Schwartz L, Gordon LI, Advani R, Jacene HA, et al. Refinement of the Lugano Classification lymphoma response criteria in the era of immunomodulatory therapy. Blood. 2016;128(21):2489–96.

23. Kurtz DM, Green MR, Bratman SV, Scherer F, Liu CL, Kunder CA, et al. Noninvasive monitoring of diffuse large B-cell lymphoma by immunoglobulin high-throughput sequencing. Blood. 2015;125(24):3679–87.

24. Daigle S, McDonald AA, Morschhauser F, Salles G, Ribrag V, McKay P, et al. Discovery of Candidate Predictors of Response to Tazemetostat in Diffuse Large B-Cell Lymphoma and Follicular Lymphoma Using NGS Technology on ctDNA Samples Collected Pre-Treatment. Blood. 2017;130(Suppl 1):4013-.

25. Sakata-Yanagimoto M, Nakamoto-Matsubara R, Komori D, Nguyen TB, Hattori K, Nanmoku T, et al. Detection of the circulating tumor DNAs in angioimmunoblastic T- cell lymphoma. Ann Hematol. 2017;96(9):1471–5.

26. Herrera AF, and Armand P. Minimal Residual Disease Assessment in Lymphoma: Methods and Applications. J Clin Oncol. 2017;35(34):3877–87.

27. Scherer F, Kurtz DM, Diehn M, and Alizadeh AA. High-throughput sequencing for noninvasive disease detection in hematologic malignancies. Blood. 2017;130(4):440–52.

28. Genovese G, Kahler AK, Handsaker RE, Lindberg J, Rose SA, Bakhoum SF, et al. Clonal hematopoiesis and blood-cancer risk inferred from blood DNA sequence. N Engl J Med. 2014;371(26):2477–87.

29. Jaiswal S, Fontanillas P, Flannick J, Manning A, Grauman PV, Mar BG, et al. Age-related clonal hematopoiesis associated with adverse outcomes. N Engl J Med. 2014;371(26):2488–98.

30. Razavi P, Li BT, Brown DN, Jung B, Hubbell E, Shen R, et al. High-intensity sequencing reveals the sources of plasma circulating cell-free DNA variants. Nat Med. 2019;25(12):1928–37.

31. Jiang L, Gu ZH, Yan ZX, Zhao X, Xie YY, Zhang ZG, et al. Exome sequencing identifies somatic mutations of DDX3X in natural killer/T-cell lymphoma. Nat Genet. 2015;47(9):1061–6.

32. Calo E, and Wysocka J. Modification of enhancer chromatin: what, how, and why? Mol Cell. 2013;49(5):825–37.

33. Chen C, Liu Y, Rappaport AR, Kitzing T, Schultz N, Zhao Z, et al. MLL3 is a haploinsufficient 7q tumor suppressor in acute myeloid leukemia. Cancer Cell. 2014;25(5):652–65.

34. Ortega-Molina A, Boss IW, Canela A, Pan H, Jiang Y, Zhao C, et al. The histone lysine methyltransferase KMT2D sustains a gene expression program that represses B cell lymphoma development. Nat Med. 2015;21(10):1199–208.

35. Lv S, Ji L, Chen B, Liu S, Lei C, Liu X, et al. Histone methyltransferase KMT2D sustains prostate carcinogenesis and metastasis via epigenetically activating LIFR and KLF4. Oncogene. 2018;37(10):1354–68.

