## Supplemental table 1 for "Genomic profiling of plasma circulating tumor DNA reveals genetics and residual disease in extranodal NK/T-cell lymphoma"

| No. | APC | ARID1A | ARID1B | ARID2 | ASXL3 | ATM | CD28 | CHD8 | CREBBP | IDH2 | JAK1 | JAK3 | KDM6A | KMT2D | NF1 | NOTCH | RHOA | SETD2 | STAT3 | TET1 | TET2 | TNFRSF | TP53 | TRAF3 |
|---|---|---|---|---|---|---|---|---|---|---|---|---|---|---|---|---|---|---|---|---|---|---|---|---|
| 1 | | | | 0.05 | | 0.06 | | 0.06 | | | | 0.05 | 0.10 | | | 0.06 | | | | | | | | |
| 2 | | | | | | | | | 0.35 | | | | | | | | | | | | | | | |
| 3 | | | | | | | | | | | | 0.20 | | | | | | | | | | | | |
| 4 | 0.10 | | | 0.19 | | | | | | | | 0.24 | | 0.18 | | | | | 0.24 | | | | | |
| 5 | | | | | | | | | | 0.47 | | | | | | 0.50 | | | | | | | | |
| 6 | | | | | | | | | | | | | | | | | | | | | | | | |
| 7 | | | | | | | | | | | | | | | | | | | | | | | | |
| 8 | | | | | | | 0.51 | | | | | | | | | | | | | | | | | |
| 9 | | | | | | | | | 0.35 | | | | | | | | | | | | | | | |
| 10 | | | | | | | | 0.49 | | | | | | | | | | | | | | | | |
| 11 | 0.47 | | | | | | | | | | | | | | | 0.55 | | | | 0.49 | | | | |
| 12 | | | | | | | | | | | | | | 0.48 | | | | | | | | | | |
| 13 | | | 0.56 | | | 0.36 | | | | | 0.56 | | | 0.50 | | | 0.15 | | | | 0.17 | | 0.61 | 0.65 |
| 14 | | | | | | | | | | | | | | 0.58 | | | | | | | | | | |
| 15 | | | | | | | | | | | | | | | | | | | | | | | | |
| 16 | | | | | | 0.44 | | | | | | | | | | | | | | | | | | |
| 17 | 0.65 | | | | | | | | | | | | | | | | | | | | | | | |
| 18 | 0.57 | | | | | | | | | | | | | | 0.48 | | | | | | | 0.54 | | |
| 19 | | | | | | | | | | | | | | | | | | | | | | | | |
| 20 | | | | | | | | | | | | | | | | | | | | | | | | |
| 21 | | | | | | | | | | | | | | | | | | | | | | | 0.61 | |
| 22 | | | | | | | | | | | | | | 0.54 | | | | | | | | | | |
| 23 | | | | | | | | | | | | | | | | | | 0.39 | | | | | | |
| 24 | | | | | 0.96 | | | | | | | | | 0.55 | | | | 0.48 | | | 0.52 | | 0.49 | |
| 25 | | | | | | | | | | | | 0.33 | | 0.22 | | | | | | | | | | |
| 26 | | | | | 0.37 | | | | | | | | | | | | | | | | | | | |
| 27 | | | | | 0.37 | | | | | | | | | | | | | | | | | | | |
| 28 | | | | | | 0.67 | | | | | | | | | | | | | | | | | | |
| 29 | | | | | | | | | | | | | | | | | | 0.43 | | | | | | |
| 30 | | | | | | | | | | | | | | | | | | | | | | | | |
| 31 | | | | | | | | | | | | | | | | | | | | | | | | |
| 32 | 0.30 | | | | | | | | | | | | | | | | | | | | | | | |
| 33 | | | | | | | | | | | | | | 0.22 | | | | | | | | | | |
| 34 | | | | | | | | | | | | | | | | | | | | | | | | |
| 35 | | 0.58 | | | | | | | | | | | | | | | | | | | | | | |
| 36 | | | | | | 0.53 | | | | | | | | | | | | | | 0.42 | | | | |
| 37 | | | | | | | | | | | | | | | | | | | | | 0.46 | | | |
| 38 | | | | | | | | | | | | | | | | | | | | | 0.53 | | | |
| 39 | 0.10 | | | 0.19 | | | | | | | | 0.24 | | 0.18 | | | | | | | | | | |
| 40 | | | | | | | | | | 0.47 | | | | | | 0.50 | | | | | | | | |
| 41 | | | | | | | | | | | | | | | | | | | | | | | | |
| 42 | | | | | | | | | | | | | | | | | | | | | | | | |
| 43 | | | | | | | 0.51 | | | | | | | | | | | | | | | | | |

| | | | | | | | | | | | | | | | | | | | | | | | | |
|---|---|---|---|---|---|---|---|---|---|---|---|---|---|---|---|---|---|---|---|---|---|---|---|---|
| 44 | | | | | | | | | 0.35 | | | | | | | | | | | | | | | |
| 45 | | | | | | | | | | | | | | | | | | | | | | | | |
| 46 | | | | | | | | | | | | | | | | 0.55 | | | | 0.49 | | | | |
| 47 | | | | | | | | | | | | | | 0.48 | | | | | | | | | | |
| 48 | | | 0.56 | | | 0.36 | | | | | 0.56 | | | 0.50 | | | 0.15 | | | | 0.17 | | 0.61 | 0.65 |
| 49 | | | | | | | | | | | | | | 0.58 | | | | | | | | | | |
| 50 | | | | | | | | | | | | | | | | | | | | | | | | |
| 51 | | | | | | | | | | | | | | | | | | | | | | | | |
| 52 | 0.65 | | | | | | | | | | | | | | | | | | | | | | | |
| 53 | 0.57 | | | | | | | | | | | | | | 0.48 | | | | | | | 0.54 | | |
| 54 | | | | | | | | | | | | | | | | | | | | | | | | |
| 55 | | | | | | | | | | | | | | | | | | | | | | | | |
| 56 | | | | | | | | | | | | | | | | | | | | | | | 0.61 | |
| 57 | | | | | | | | | | | | | | 0.54 | | | | | | | | | | |
| 58 | | | | | | | | | | | | | | | | | | 0.39 | | | | | | |
| 59 | | | | | 0.96 | | | | | | | | | 0.55 | | | | 0.48 | | | | | 0.49 | |
| 60 | | | | | | | | | | | | 0.33 | | 0.22 | | | | | | | | | | |
| 61 | | | | | 0.37 | | | | | | | | | | | | | | | | | | | |
| 62 | | | | | 0.37 | | | | | | | | | | | | | | | | | | | |
| 63 | | | | | | 0.67 | | | | | | | | | | | | | | | | | | |
| 64 | | | | | | | | | | | | | | | | | | 0.43 | | | | | | |
| 65 | | | | | | | | | | | | | | | | | | | | | | | | |
